# Decomposing Loss Aversion from Gaze Allocation and Pupil Dilation

**DOI:** 10.1101/2020.02.27.967711

**Authors:** Feng Sheng, Arjun Ramakrishnan, Darsol Seok, Wenjia Joyce Zhao, Samuel Thelaus, Puti Cen, Michael Louis Platt

**Author notes:** Correspondence should be addressed to (F.S.), (A.R.), or (M.L.P.). F.S and A.R. contributed equally. D.S. and W.J.Z. contributed equally.

## Abstract

Loss-averse decisions, in which one avoids losses at the expense of gains, are highly prevalent. However, the underlying mechanisms remain controversial. The prevailing account highlights a valuation bias that overweighs losses relative to gains, but an alternative view stresses a response bias to avoid choices involving potential losses. Here we couple a computational process model with eye-tracking and pupillometry to develop a physiologicallygrounded framework for the decision process leading to accepting or rejecting gambles with equal odds of winning and losing money. Overall, loss-averse decisions were accompanied by preferential gaze towards losses and increased pupil dilation for accepting gambles. Using our model, we found gaze allocation selectively indexed valuation bias, and pupil dilation selectively indexed response bias. Finally, we demonstrated that our computational model and physiological biomarkers can identify distinct types of loss-averse decision-makers who would otherwise be indistinguishable using conventional approaches. Our study provides an integrative framework for the cognitive processes that drive loss-averse decisions and highlights the biological heterogeneity of loss aversion across individuals.

**Significance Statement:** We revisit the concept of loss aversion by synthesizing distinct views into an integrative framework and by probing physiological biomarkers associated with the behavior. The framework decomposes loss aversion into a valuation bias, which weighs losses over gains, and a response bias, which avoids loss-related choices altogether. Further, we revealed a double dissociation in physiology underlying the decision process. Valuation bias was associated with preferential gaze allocation to losses whereas response bias was associated with pupillary dilation. Our framework exposed biological heterogeneity underlying loss aversion and distinguishes different loss-averse decision makers who are otherwise indistinguishable using conventional approaches. Our integrative approach provides a deeper analysis of the mechanisms underlying loss aversion and incorporates distinct views within a unified biological framework.

## Introduction

Many decisions involve tradeoffs between potential gains and losses. A widespread anomaly in decision making is loss aversion—the tendency to avoid losses at the expense of acquiring equivalent gains (1). For example, people often reject gambles of equal odds to gain and lose money, unless the amounts of potential gains are sufficiently larger than potential losses (2–4). Loss aversion has been observed in a variety of laboratory settings as well as in the field (see ref. 5 for a review) and is even evident in other species like chimpanzees (6), monkeys (7) and rats (8). Despite the ubiquity of loss aversion, the underlying mechanisms remain unresolved and hotly debated.

Prospect theory attributes loss-averse decisions to a *valuation bias* that overweighs losses relative to gains (1, 9). According to this view, a chooser would require an extra amount of potential gain, known as a premium, to accept a mixed gamble, in order to compensate for the lower weight of gain relative to loss. Decision making, however, is not always based on valuation alone. For example, it is robustly observed that preference ratings and choices are not always aligned (see (10) for a review). In fact, decision making is dramatically susceptible to context (10, 11), including the possible responses available (12–16). Accordingly, an alternative account attributes loss-averse decisions to a *response bias* that inclines choosers to avoid the choice or action that may incur losses (14, 17), and remain with the status quo (18). According to this view, choosers have a predisposition to avoid the choice that can change the status quo, by rejecting a gamble, rather than endure the possibility of losing, by accepting a gamble. Thus, even when potential gains and losses are weighted equally, a premium is required to overcome the resistance to accept the gamble. These two accounts are often indistinguishable based solely on the decisions people make.

Most decisions, loss-averse or otherwise, are not made instantaneously. Abundant evidence indicates that many decisions unfold through a process of evidence accumulation over time that culminates in choice (19, 20). Evidence-accumulation is evident in neurophysiological findings in humans (21), monkeys (22) and rats (23), and is formalized in computational process models like the drift-diffusion model (DDM) (20). Although valuation bias and response bias can result in the same decision, in theory they shape the evidence accumulation process in distinct ways. Specifically, valuation bias would impact how information (e.g., potential gain and loss) is evaluated over time, whereas response bias would impact how much evidence must be accumulated to make a decision (e.g., accept vs. reject a gamble). Thus, examining the decision process as it unfolds could potentially allow us to disentangle the two biases.

Recent studies have attempted to understand the decision process and examined the strengths of these biases in loss-averse decisions by simultaneously fitting DDM models to decisions and the time it takes to make them (24, 25). Response times, however, provide at best an indirect window on the internal deliberative process. To overcome this limitation, here we coupled a computational process model with eye-tracking and pupillometry to provide a physiologicallygrounded framework for the decision process underlying loss-averse decisions.

Where we look betrays our underlying biases and predicts choice (26). As a selective filter on information acquisition, gaze dynamics reveal the otherwise hidden process of evidence accumulation (27). Specifically, gaze reflects and amplifies the subjective value of fixated options (27, 28) or features (29), and thereby shapes the decision process. We hypothesized that valuation bias, which overweighs losses relative to gains as evidence supporting choice, would manifest as biased gaze toward losses over gains during deliberation.

Pupil dilation, in the absence of changes in external lighting, indexes activation of the arousal system in the brain (30, 31). Higher arousal and pupil dilation accompany effortful decisions (31, 32). Making an infrequent choice that contradicts a more habitual one requires effort to overcome inertia, and evokes pupil dilation (33). Within the DDM framework, larger pupil dilation during decision making reflects a higher response threshold for making a decision (28). We hypothesized that if there is a default response bias to avoid an option that may incur losses, actually choosing that option should require more effort and therefore evoke larger pupil dilation.

In summary, we propose that valuation bias and response bias can be simultaneously accommodated within the framework of an evidence accumulation-based decision process. Specifically, gaze allocation would reflect valuation bias whereas pupil dilation would reflect response bias. To test our hypotheses, we examined the decision process underlying choices for mixed gambles while tracking gaze and measuring pupil diameter. Participants (N=94) chose to accept or reject a series of gambles offering a 50% chance of winning or losing money ranging from $1 to $10, which were displayed on a computer screen (see Fig. 1A and Materials and Methods for more details).

**Fig. 1.**
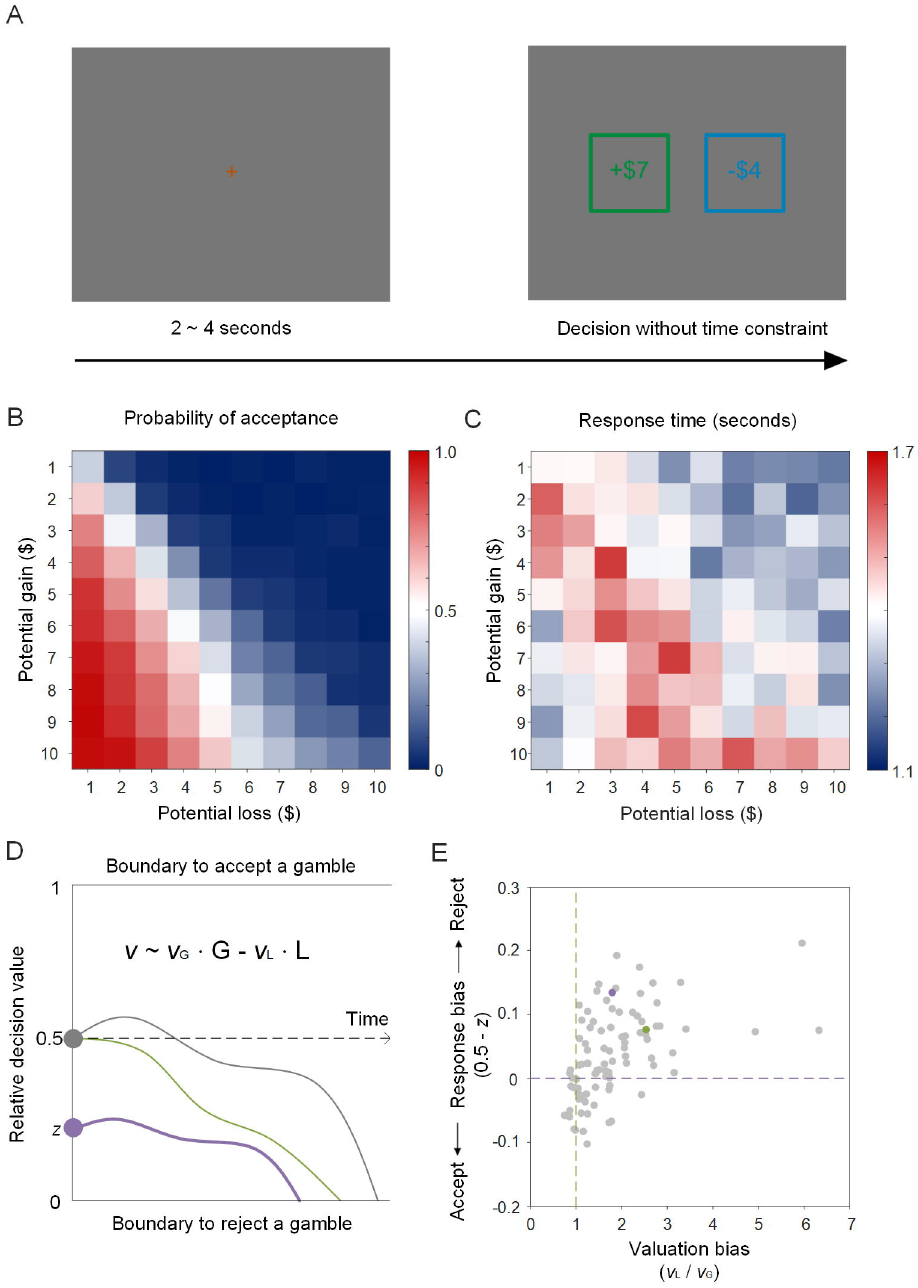
Behavior. Top panel (*A*): gambling task. Gambles were not resolved after decisions. Middle panel: probability of acceptance (*B*) and response times (*C*) across gambles. Bottom panel: illustration of the drift-diffusion model for the gambling task (*D*) and estimates of valuation bias and response bias for each individual participant (*E*). Each dot represents a participant: those to the right of the green dashed line displayed valuation bias of overweighing loss relative to gains, and those above the purple dashed line displayed a response bias to reject gambles. The green dot (P69) and the purple dot (P46) in quadrant 1 indicate the two example participants illustrated in ***SI Appendix,* Fig. S6** and **Fig. S7**.

## Results

Participants in our study showed a typical pattern of loss aversion in their choices (Fig. 1B) and response times (Fig. 1C) (2–4, 24). Despite our symmetric design of gains and losses across gambles, overall participants were less likely to accept than reject gambles *(SI Appendix,* Fig. S1A, M±SD=31±16%, ranging from 4% to 78%, t(93)=-11.357, p<0.001) and were slower to accept *(SI Appendix,* Fig. S1B, M±SD=1.595±0.440 s, t(93)=-6.195, p<0.001) than reject (M±SD=1.369±0.405 s) gambles. Moreover, they demanded a premium over an expected value of zero to be indifferent to accepting or rejecting gambles (*SI Appendix,* Fig. S1C, M±SD=$1.266±1.216, t(93)=10.098, p<0.001; also see white cells in Fig. 1B) and they took the longest time to make decisions for gambles entailing such a premium (*SI Appendix,* Fig. S1D, M±SD=$1.271±2.269, t(93)=5.432, p<0.001; also see red cells in Fig. 1C).

**Fig. 2.**
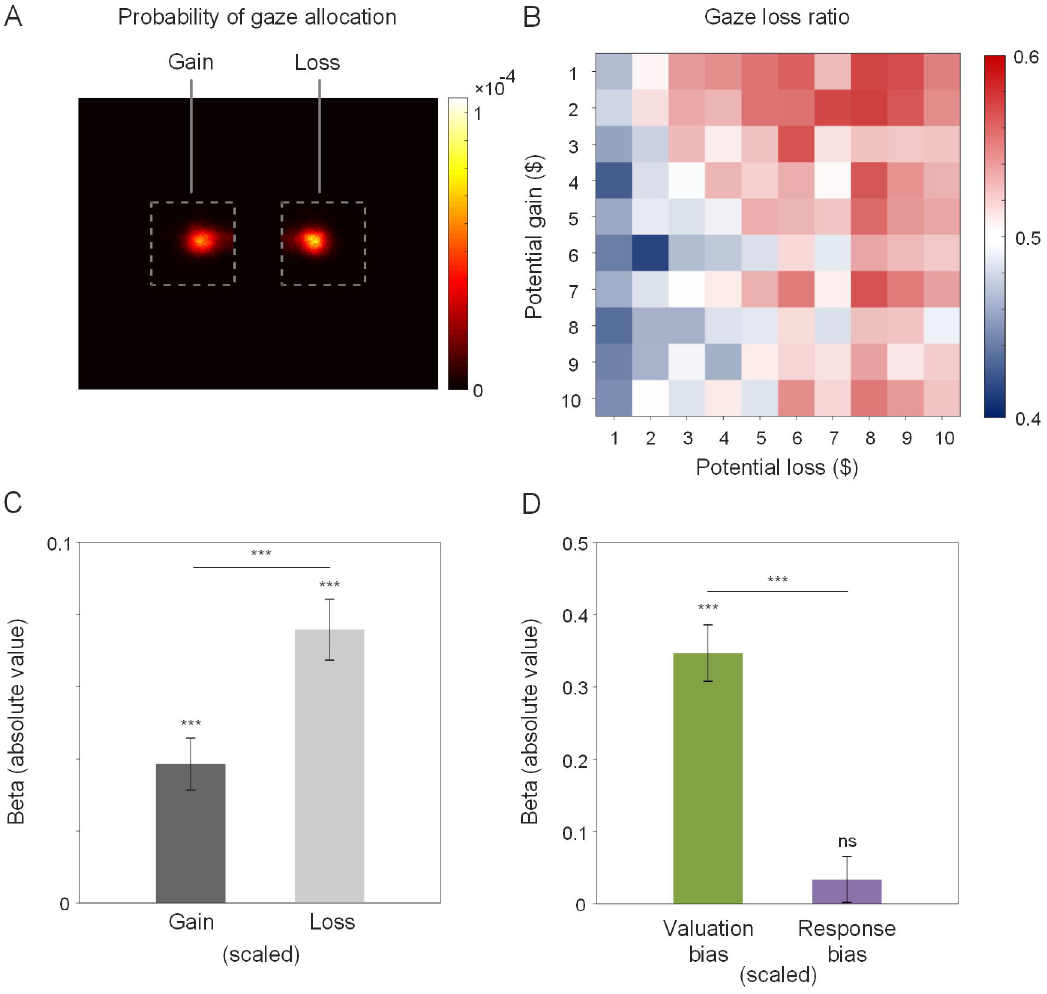
Gaze. (*A*) Probability of gaze allocation to gains and losses. (*B*) Gaze-loss ratio across gambles. (*C*) Gaze-loss ratio predicted by magnitude of gain and loss across trials. Black and gray bars denote negative and positive beta values, respectively. (*D*) Gaze-loss ratio predicted by valuation bias and response bias across participants. Error bars indicate standard errors. ***: p<0.001; ns: nonsignificant, p>0.1.

To provide a simultaneous account of choice probabilities and response times with valuation bias and response bias, we adapted the DDM from ref. 24. In the model, each decision was modeled as the outcome of a noisy evidence accumulation process reaching one of two response boundaries, either accepting (upper, 1) or rejecting (lower, 0) the gamble (Fig. 1D). The speed of evidence accumulation, or drift rate (*v*), depended on the amounts of gain *(G)* and loss *(L)* weighted by their respective coefficients (*ν_G_* and *ν_L_*). That is, *v ~ ν_G_* · *G* – *ν_L_* · *L*. In this framework, the ratio between the two coefficients (*ν_l_/ν_G_*) represented valuation bias. Response bias was captured by the starting point *(z)* of the evidence accumulation process. A starting point in the middle of the two response boundaries (*z*=0.5) indicated no *a priori* response bias, whereas a starting point closer to the rejection boundary indicated a response bias toward rejecting the gamble (quantified as 0.5-*z*). According to this model, a person is more likely and/or faster to reject a gamble than another person either due to a valuation bias—manifested as a steeper drift rate toward the rejection boundary (e.g., green vs. gray curve in Fig. 1D)—or due to a response bias—manifested as a starting point closer to the rejection boundary (e.g., purple vs. gray curve, in Fig. 1D). By fitting this model simultaneously with choice probabilities and response times using a hierarchical Bayesian approach (see Materials and Methods for more details), we were able to dissociate and quantify valuation bias and response bias for each individual participant (Fig. 1E).

Both valuation bias (group mean=1.575, 95% Credible Interval: 1.398 to 1.780,greater than 99.9% likelihood > 1) and response bias (group mean=0.037, 95% Credible Interval: 0.023 to 0.052, greater than 99.9% likelihood > 0) were robustly present in the population. Across individuals, valuation bias ranged from strong overweighting of loss to mild underweighting of loss (Fig. 1E, 6.323 to 0.749) and response bias ranged from a strong tendency to reject gambles to a mild tendency to accept gambles (0.211 to −0.103), indicating heterogeneity across participants. The correlation between the two biases was moderate (r(92)=0.488, p<0.001), suggesting that valuation bias and response bias may be related but do not necessarily index the same internal process. To further illustrate the independence of the two biases, we formally compared our model, which incorporated both valuation bias and response bias (i.e. *ν_G_, ν_L_*, and *z* were all free parameters), with three reduced models: one that incorporated only valuation bias but no response bias (i.e., *z*=0.5); a second that incorporated only response bias but no valuation bias (i.e., *ν_G_=ν_L_*); and a third model that left out both biases (*ν_G_=ν_L_* and *z*=0.5). We found that the model that incorporated both biases best explained the variance in choices and response times (see *SI Appendix,* Fig. S2 and Table S1). Thus, both valuation bias and response bias are necessary to explain the observed behaviors in our study.

Next, we examined the relationship between valuation bias and gaze allocation, and response bias and pupil dilation. On average, before a decision was made, participants made 2.710±1.434 gaze fixations on the displayed gain and loss amounts (median=3, *SI Appendix,* Fig. S3). A heatmap plot of gaze allocation probability suggested an overall tendency to inspect information about loss relative to gain (Fig. 2A). To quantify potential gaze bias, we calculated *gaze-loss ratio* for each trial as the time spent looking at the loss amount relative to the total time spent looking at both gain and loss. We also calculated participant-level gaze-loss ratio by averaging gaze-loss ratios across trials for each individual. We found that gaze-loss ratio was slightly, but not significantly, larger than 0.5 in our population (M±SD=0.513±0.085, range from 0.389 to 0.904, one-sample t-test against 0.5, t(92)=1.428, p=0.078), indicating heterogeneity in gaze allocation to gains and losses across the population, despite an overall tendency to attend more to losses.

Notably, gaze-loss ratio was sensitive to the magnitudes of gain and loss (Fig. 2B). A multi-level regression revealed that gaze-loss ratio increased as the magnitude of potential loss increased (Fig. 2C, B=0.076, SE=0.008, t(17425)=8.97, 95% Confidence Interval: 0.059 to 0.092, p<0.001) and decreased as the magnitude of potential gain increased (B=-0.038, SE=0.007, t(17425)=5.34, 95% Confidence Interval: −0.053 to −0.024, p<0.001). Importantly, increases in the magnitude of loss had nearly double (1.966 times) the impact of increases in the magnitude of gain on gaze-loss ratio (Fig. 2C, linear contrast of betas, F(1,17425)=15.099, p<0.001), an asymmetric pattern (Fig. 2B) similar to that observed in choices (Fig. 1B) and response times (Fig. 1C). These results suggest that gaze allocation is attracted by loss during information sampling, possibly reflecting valuation bias in the underlying evidence accumulation process.

To test this hypothesis, we used regression analysis to determine whether valuation bias and response bias estimated from DDM predicted individual differences in gaze-loss ratio. We found that valuation bias (Fig. 2D, B=0.347, SE=0.039, t(90)=8.894, 95% Confidence Interval: 0.270 to 0.425, p<0.001) but not response bias (B=0.034, SE=0.032, t(90)=1.063, 95% Confidence Interval: −0.029 to 0.097, p=0.291) predicted gaze-loss ratio. The predictive power of valuation bias on gaze-loss ratio was 10.284 times larger than that of response bias (linear contrast of betas, F(1, 90)=26.28, p<0.001). Consequently, our findings endorse a strong relationship between gaze allocation and valuation bias.

We then tested our hypothesis that pupil dilation reflects response bias. We predicted that accepting a gamble would be effortful and would therefore evoke pupil dilation in choosers who had a default bias to reject gambles offering potential loss. As expected, we observed that pupil size began to increase about 0.5 s before accepting a gamble, and, as a sluggish physiological response, pupil size continued to increase for about 1.5 s after the decision (Fig. 3A, *SI Appendix*, Fig. S4A), a time course consistent with earlier research on pupil dilation and decision making (33). We calculated the mean pupil size in this time window ranging from −0.5 s before to 1.5 s after decision as *decision-related pupil size.* Overall, participants showed larger decision-related pupil size when accepting (M±SD=0.151±0.411) than when rejecting gambles (M±SD= −0.088±0.314, paired t-test, t(92)=8.111, p<0.001). Moreover, participants who accepted fewer gambles showed larger pupil dilation when they accepted gambles compared to when they rejected them (Fig. 3B, r(91)=-0.661, p<0.001). This correlation was primarily driven by gamble acceptance (*SI Appendix,* Fig. S4B, red dots and line, r(91)=-0.417, p<0.001) but not by gamble rejection (blue dots and line, r(91)=0.053, p=0.617). These observations suggest that accepting a gamble or rejecting it were asymmetric decisions for participants.

**Fig. 3.**
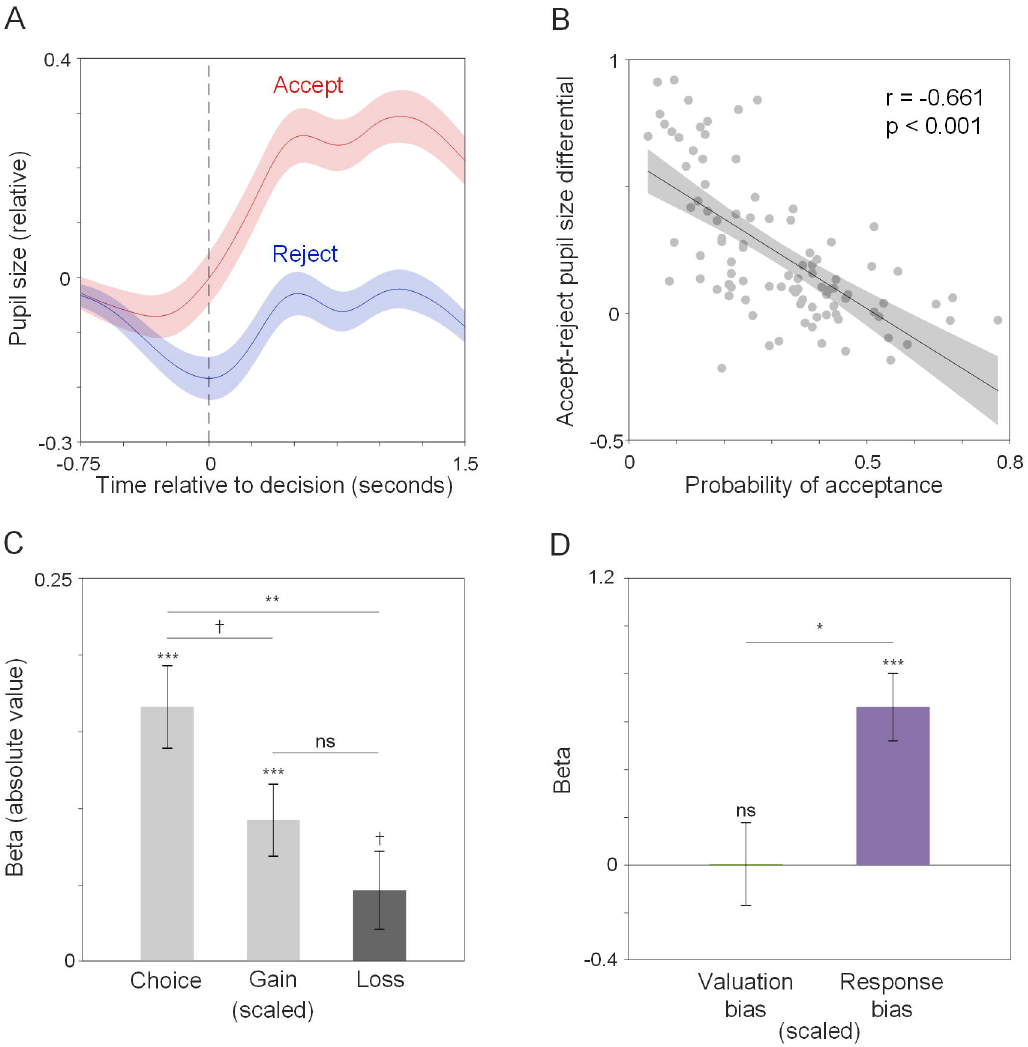
Pupil. *(A)* Change in pupil size around the time of the decision (t=0) to accept (red) vs. reject (blue) gambles. *(B)* Correlation between probability of gamble acceptance and accept-reject pupil size differential. *(C)* Influence of choice, gain and loss on decision-related pupil size across trials. Black and gray bars denote negative and positive beta values, respectively. *(D)* Accept-reject pupil size differential predicted by valuation bias and response bias. Error bars indicate standard errors. ***: p<0.001; **: p<0.01; *: p<0.05; †: p<0.1; ns: nonsignificant, p>0.1.

Gambles accepted, relative to those rejected, usually entailed larger gains and smaller losses (Fig. 1B), or larger expected values (*SI Appendix,* Fig. S1C). Thus, the observed difference in decision-related pupil size of accepted vs. rejected gambles, which we refer to as *accept-reject pupil size differential,* might simply reflect differences in the expected value of gambles, rather than a differential response to the choice itself. To unveil the influence of choice itself on pupil dilation, we performed a mixed-level regression to predict decision-related pupil size with gain, loss and choice (accept=1, reject=0). We found that choice was in fact the biggest contributor to decision-related pupil size (Fig. 3C, B=0.166, SE=0.027, t(17424)=6.194, 95% Confidence Interval: 0.114 to 0.219, p<0.001), compared to gain (B=0.092, SE=0.023, t(17424)=3.935, 95% Confidence Interval: 0.046 to 0.138, p<0.001) or loss (B=-0.047, SE=0.025, t(17424)=1.841, 95% Confidence Interval: −0.097 to 0.003, p=0.066). The influence of choice on decision-related pupil size was 1.798 times greater than that of gain (linear contrast of betas, F(1,17424)=3.623, p=0.057) and 3.550 times greater than that of loss (linear contrast of betas, F(1,17424)=8.406, p=0.004).

These results indicate that decision-related pupil size is primarily influenced by the decision itself rather than the values of the gambles, and likely reflects response bias. To test this idea, we examined whether response bias and valuation bias estimated from the DDM predicted accept-reject pupil size differential using regression. We found that larger response bias (Fig. 3D, B=0.661, SE=0.141, t(90)=4.706, 95% Confidence Interval: 0.382 to 0.940, p<0.001) but not valuation bias (B=0.005, SE=0.173, t(90)=0.029, 95% Confidence Interval: −0.338 to 0.348, p=0.977) predicted larger accept-reject pupil size differential across individuals. The effect of response bias was significantly larger than that of valuation bias (linear contrast of betas, F(1,90)=5.882, p=0.017). These findings support the idea that decision-related pupil size strongly reflects response bias when considering whether to accept or reject a gamble.

## Discussion

By considering decision making as an evidence accumulation process that unfolds over time, the present study synthesizes two distinct views of loss aversion into a single, biologically-grounded framework. Specifically, by fitting DDM to choice probabilities and response times observed in decisions over mixed gambles, we quantitatively decomposed loss aversion into a valuation bias that weighs loss over gain and a response bias that promotes gamble avoidance prior to the evaluation of gain and loss amounts. Our approach revealed heterogeneity in the mechanisms underlying loss aversion (Fig. 1E), which would be otherwise undetectable using conventional approaches like prospect theory (PT) that do not explicitly incorporate both biases (3, 9).

In our sample, loss aversion, when estimated as a single bias by PT (denoted as λ, see Materials and Methods for details), was correlated with both valuation bias (r(92)=0.343, p<0.001) and response bias (r(92)=0.588, p<0.001) obtained from DDM across individuals. This suggests that PT-estimated loss aversion shares variability with both valuation bias and response bias. To better understand the relationship between PT and DDM, we plotted the ratio of PT-estimated loss aversion to valuation bias (i.e., λ/(*ν_L_/ν_G_*)) against response bias (*SI Appendix,* Fig. S5). This ratio was larger than 1 for more participants (57%) when response bias was positive (i.e., quadrant 1 vs. 4), but smaller than 1 for more participants (62%) when response bias was negative (i.e., quadrant 3 vs. 2), which resulted in a positive correlation between the ratio and response bias (r(92)=0.207, p=0.045). That is, loss aversion estimated by PT tended to be larger than valuation bias when there was a synergistic response bias toward rejecting gambles, and smaller when there was a counteractive response bias toward accepting gambles. Thus, loss aversion estimated by PT reflects the combined effects of both response bias and valuation bias.

By decomposing loss aversion into two components, our DDM framework helps discriminate loss-averse decision makers in a more nuanced way. Two individuals who are similarly loss-averse according to PT can be distinct in response bias and valuation bias in our DDM framework. For example, in our sample, P69 (the green dot in Fig. 1E and *SI Appendix,* Fig. S5) and P46 (the purple dot) accepted similar sets of gambles (probability of acceptance: 22.5% vs. 23.0%, *SI Appendix,* Fig. S6A & B). As a result, loss aversion estimated by PT, which only modeled choices and assumed no *a priori* response bias, was roughly equivalent for these two participants (λ: 2.16 vs. 1.98). In our DDM-based approach, which takes into account response times (*SI Appendix,* Fig. S6C & D) in addition to choices, P46 showed a larger response bias than did P69 (0.13 vs. 0.08), whereas P69 evinced a larger valuation bias than did P46 (2.54 vs. 1.79, see Fig. 1E and *SI Appendix,* Fig. S5).

The DDM framework also accommodates instances when valuation bias and response bias counteract one another (Fig. 1E, quadrant 2 & 4). For example, in our sample, a non-negligible portion of participants (20.21%) showed valuation bias for loss over gain but also showed a ‘reverse’ response bias favoring gamble acceptance (Fig. 1E, quadrant 4). For these participants, loss aversion estimated using PT would be a compromise of the two biases. These cases may partly explain why loss aversion was not observed in some prior studies or why observed loss aversion was sometimes vulnerable to response framing (17, 34, 35).

By using eye-tracking and pupil monitoring to unmask the internal deliberative process, we found that valuation bias and response bias are indexed by distinct physiological processes. Specifically, gaze allocation selectively reflected valuation bias and pupil dilation selectively reflected response bias. This double dissociation in physiology further validates our approach to decomposing loss aversion. Valuation bias and response bias are not only mathematically distinguishable through application of DDM (24), but are also biologically separable. This finding provides biomarkers for us to distinguish distinct types of loss-averse decision makers even when their choices were similar. For instance, among the two example participants, P69 (the green dot in Fig. 1E and *SI Appendix,* Fig. S5), who showed higher valuation bias, also displayed a larger gaze-loss ratio (59.98% vs. 52.61%, *SI Appendix,* Fig. S7A & B). Conversely, P46 (the purple dot in Fig. 1E and *SI Appendix,* Fig. S5), who showed a higher response bias, also evinced larger accept-reject pupil size differential (0.804 vs. 0.392, *SI Appendix,* Fig. S7C & D).

The role of visual attention in loss-averse decisions was tested indirectly in a recent study that tracked mouse-movements in computer-based gambling tasks (36). The present study, by coupling eye-tracking with DDM, provides direct evidence for a gaze bias toward loss involved in the evidence accumulation process that culminates in loss-averse decisions. A potential mechanism for gaze to impact this process is by dynamically altering the weights of fixated vs. non-fixated information during evidence accumulation, as proposed by the attentional drift-diffusion model (aDDM) (27). Applying the aDDM to our data (see Materials and Methods for details) showed that valuation bias only partially depends on gaze. Information about loss was weighted more heavily than information about gain regardless of whether gaze was on gain or loss. Importantly, the overweighting of loss relative to gain manifested to an even larger degree when gaze was on loss than when it was on gain (see *SI Appendix,* Supplementary Text and Fig. S8 for details). Based on these findings, one possible mechanism is that loss-averse decision makers have an *a priori* inclination to weigh loss over gain, which drives them to preferentially inspect information about loss relative to gain by looking at it, and this gaze bias, in turn, further enhances the weight of loss in the evidence accumulation process (also see ref. 26). Future studies, particularly causal manipulations (e.g., ref. 36), will be necessary to thoroughly address the role of gaze and its influence on valuation and decision making.

In binary decisions without *a priori* bias, increase in pupil dilation reflects increase in the decision threshold for choices in the DDM framework (28). The present work extends this view by showing that when there is an *a priori* bias for one choice (e.g., rejecting a gamble), larger pupil dilation is evoked by choosing the alternative (e.g., accepting the gamble), which is captured by a biased starting point in the DDM framework. In both cases, larger pupil dilation represents a longer journey traversed by the evidence accumulation process.

In the present study, we observed pupil responses began to depart about 500 ms before an upcoming choice to accept vs. reject a gamble and the trend continued after the decision. Pupil dilation is a sluggish physiological signal, which is often delayed and prolonged with respect to its cause (33, 37). Thus, the underlying decision process for accepting vs. rejecting a gamble may have begun to diverge even earlier.

Based on these findings, a potential mechanism underlying the decision to accept a gamble, especially in individuals with large response bias, could be the following: After gamble presentation, gaze allocation on potential gain may lead to reevaluation of the default bias to reject a gamble. This reevaluation could slow down the decision process, leading to an increase in response time, and generate efforts to update belief and implement the non-default choice to accept the gamble, which would ultimately be manifested as pupil dilation.

Our decomposition approach also provides a framework to organize the mixed neurophysiological findings about loss aversion. While functional neuroimaging research shows larger sensitivity of the prefrontal-striatal dopaminergic system to the magnitude of loss relative to gain (2, 38), pharmacological (39) and positron emission tomography studies (40) link loss aversion to the noradrenergic arousal system, which mediates non-luminance-evoked pupil dilation (30, 31). Our findings suggest that activation of the noradrenergic system, which is indexed by changes in pupil size, may reflect the process of overcoming response bias, while the dopaminergic system may be recruited to encode the value of gain and loss at different scales (2), thus giving rise to valuation bias.

Overall, our study decomposed loss aversion into a valuation bias and a response bias, which were reflected in gaze allocation and pupil dilation, respectively. The decomposition into distinct biases offers a more nuanced, biologically-grounded perspective on the mechanisms mediating decision making and advocates for physiologically-based approaches to determine underlying biases. Our study reveals individual differences in the neurobiological mechanisms underlying loss aversion, which can lead to better understanding and personalized interventions to overcome maladaptive biases in decision making.

## Materials and Methods

### Participants

The study was approved by the ethics committee of the University of Pennsylvania. Ninety-four adults (54 females, 38 males, 2 non-identified; 18-56 years old, M±SD=22.82±6.39) participated in the experiment in three cohorts. Results reported here were based on all three cohorts of participants and similar results were found for each single cohort. All participants had normal or corrected-to-normal vision. One participant was excluded for gaze and pupil analysis because he thought he was required to maintain fixation on the center of the screen and did not allocate any fixation on gain or loss in more than 90% of the trials.

### Procedure

In the gambling task, participants decided whether to accept or reject gambles of equal chance of winning and losing money on a computer (Fig. 1A). Each trial started with a 2~4 s of gaze fixation period, followed by a gamble with the amounts of potential gain and loss displayed in two boxes positioned at the left and right to the fixation, separated by about 13 degrees. Participants were given unlimited time to make the decision by pressing one of two keys. Note, the gambles were not immediately resolved post-decision.

The amounts of potential gain and loss ranged from $1 to $10, with $1 increment. This resulted in a total of 100 unique gambles. Each participant faced two such blocks of 100-trials each. In other words, they faced each gamble twice, once in each block. The positions of the gain and loss—left vs. right of the fixation cross—were counterbalanced between blocks. For a random half of the participants, gain was displayed in blue and loss in green and for the other half the colors were switched. The fixation cross was orange in color and the background was gray. Colors were adjusted to make the screen isoluminant.

At the start of the experiment, participants received an endowment of $10 in cash. They were instructed that at the end of the experiment, one trial (out of 200) would be randomly selected and a payment made according to their actual decision. If the selected gamble was rejected, participants left with the original $10. If the gamble was accepted, a coin would be flipped to determine whether they won or lost the gamble. If they won, the amount of gain was added to the tally and if they lost, the amount of loss was subtracted from it. Thus, decisions directly impacted payments, making the task incentive-compatible.

### Gaze and Pupil Data Acquisition

Participants were seated about 60 cm in front of the screen in a dark and silent room with their head stabilized using a chin rest. Gaze fixation and pupil diameter were sampled at 120 Hz using an infra-red eye-tracker (SensoMotoric Instruments, Germany). The eye-tracker was synchronized with the stimulus-presentation software (iMotions, Denmark).

### Discarding of Eye-Tracking Data

Trials with less than 50% of the gaze or pupil data in either the analysis time window (i.e., 0.5 s before gamble onset to 1.5 s after decision) or in the baseline time window (i.e., 0.5 s to 0 s before gamble onset) were excluded, leaving 97.37% ‘valid’ trials.

### Processing of Gaze Data

The boxes containing the amounts of gain and loss on the screen were identified as regions of interests. Total time spent looking at the gain and loss regions in a trial was calculated and referred to as ‘gaze duration’ of gain or loss. Short gaze fixations (< 30 ms) were discarded for this calculation. Gaze-loss ratio was calculated for the trials that included at least one gaze fixation on either gain or loss (96.23% out of all valid trials). Heatmaps of gaze allocation probability were created by calculating the percent of gaze samples falling on each pixel within the regions of interests in a trial and then averaged across trials for each participant (*SI Appendix,* Fig. S6A & B) and the group (Fig. 2A).

### Processing of Pupil Data

Only the left pupil size was considered for analysis. Extreme or isolated pupil samples were excluded (41). Missing or excluded samples were linearly interpolated. The pupil data were then resampled to 100 Hz, filtered using a third-order Butterworth filter (low pass, 4 Hz) and z-scored for each run and each participant. For each trial, baseline correction was done by subtracting the mean value of the 0.5 s before gamble onset from each value within the analysis time window, i.e., from 0.5 s before gamble onset to 1.5 s after decision. Decision-related pupil size was calculated as the mean pupil size in the time window ranging from 0.5 s before to 1.5 s after the decision (33).

### Drift-diffusion modelDDM

A sequential sampling model was adapted from ref. 24 to account for choices and response times of the mixed gambling task. Specifically, the model assumes that the decision to accept or reject a gamble is a process of evidence accumulation, with moment-to-moment fluctuation (Fig. 1D). The accumulation process is terminated as soon as one of the two response boundaries—either accepting (upper) or rejecting (lower) the gamble is reached. The choice and the response time are jointly determined by the rate at which evidence accumulates (drift rate, *v*) and the starting point of the accumulation process (*z*), relative to the two response boundaries, i.e., 0 and 1. Valuation bias is reflected in the drift rate. For a gamble *i* with gain amount of *G_i_* and loss amount of *L_i_*, we assume *v_i_, = ν_G_* · *G_i_* – *ν_L_* · *L_i_* + *b* + *ε*. Here *ν_G_* and *ν_L_* denote coefficients or weights associated with gains and losses, respectively, and thus the ratio between the two, *ν_L_/ν_G_*, is indicative of the valuation bias. *b* is an intercept, indicating a constant component in the drift rate. *ε* indicates random noise in the diffusion process, which follows a standard normal distribution, i.e., *ε~N*(0,1). Response bias is determined by the starting point. A starting point right in the middle, or *z*=0.5, indicates no *a priori* bias toward any of the two response boundaries, and a starting point closer to the rejection boundary, or *z*<0.5, represents a response bias toward rejection, which can be quantified as 0.5-*z*.

We tested four models: DDM 1, a baseline model that constrained both valuation bias and response bias, i.e., *ν_G_=ν_L_* and *z*=0.5; DDM 2, a model that allowed response bias but constrained valuation bias, i.e. *ν_G_*=*ν_L_*; DDM 3, a model that allowed valuation bias but constrained response bias, i.e. *z*=0.5; and DDM 4, the model that enabled both valuation bias and response bias, in which, *ν_G_*, *ν_L_* and *z* were all set as free parameters. See *SI Appendix,* Table S1 for a summary. For each model, the distance of the two response boundaries (*a*) and the nondecision time (*t*) were also included as variables of non-interests.

These models were fit to choice and response time data using HDDM (42), a Python package for hierarchical Bayesian estimation of drift-diffusion models. This approach estimates group and individual level parameters simultaneously, with group-level parameters forming the prior distributions from which individual participant estimates are sampled. We ran 4 separate chains for the models. Each chain consisted of 20,000 samples, where the first 2000 were burnins. To assess model convergence, 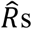 of all the parameters were calculated. All 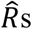 were close to 1, suggesting that the sample size was sufficient for the chains to converge (43).

To perform model comparison, we calculated Bayesian Predictive Information Criterion (BPIC) (44) for each model (*SI Appendix,* Fig. S2A). Further, we tested out-of-sample predictions of each model. Specifically, we segregated data by whether gain was presented on the left or right side of the screen on a trial. We first estimated parameters by fitting the DDMs with choice and response time data of gain-left (gain-right) trials, then used the estimated posterior means of parameters to compute log likelihood of the observed distribution of choices and response times of gain-right (gain-left) trials, and finally aggregated log likelihood across trials and participants (*SI Appendix,* Fig. S2B).

### drift-diffusion modelaDDM

To test attentional drift-diffusion model (aDDM) (27) in the framework of HDDM (also see ref. 28), we allowed the coefficients of gain and loss in the drift rate, *ν_G_* and *ν_L_*, to vary as a function of gaze. Specifically, we assume that when gaze is on gain, the coefficients for gain and loss in the drift are *ν_G,GazeG_* and *ν_L,GazeG_*, whereas when gaze is on loss, the two coefficients are *ν_G_,GazeL* and *vL,GazeL.* Thus, for a gamble *i* with gain amount of *G_i_* and loss amount of *L_i_*, the drift rate *ν_i_, ~ ν_G,Gazeo_* · *G_i_ – ν_L,GazeG_* · *L_i_* when gaze is on gain and *ν_i_ ~ ν_G,GazeL_* · *G_i_ – ν_L,GazeL_* · *L_i_* when gaze is on loss. Assuming for the gamble *i*, the percent of time spent looking at loss, or gaze-loss ratio, is *GazeL_i_* and that for gain, or gaze gain ratio, is *GazeG_i_* (i.e., 1-*GazeL_i_*), the mean drift rate for the gamble can be represented as the sum of two components weighted by respective gaze ratios, i.e., *ν_i_ ~ GazeG_i_* · (*ν_G,GazeG_* · *G_i_ – ν_L,GazeG_* · *L_i_*) + *GazeL_i_* · (*ν_G,GOzeL_* · *G_i_ – ν_L,GazeL_* · *L_i_*). Accordingly, valuation bias is *ν_L,GazeG_*/*ν_G,GazeG_* when gaze is on gain and *ν_L,GazeL_*/*ν_G,GazeL_* when gaze is on loss, and the discounting rate of being non-fixated is *ν_G,GazeL_/ν_G,GazeG_* for gain and *ν_L,GazeG_/ν_L,GazeL_* for loss.

We tested four instantiations of aDDM in this framework: aDDM 1, a baseline model that assumed no valuation bias at all, i.e., *ν_G,GazeG_ = ν_L,GazeG_* = *ν_G,GazeL_ = ν_L,GazeL_*, aDDM 2, a model that assumed valuation bias was fully dependent on gaze, i.e., *ν_G,GazeG_ = ν_L,GazeG_* and *ν_G,GazeL_ = ν_L,GazeG_*; aDDM 3, a model assumed valuation bias was fully independent of gaze, i.e., *ν_G,GazeG_ = ν_G,GazeL_* and *ν_L,GazeG_ = ν_L,GazeL_*, and aDDM 4, a model that assumed valuation bias could have both gaze-dependent and -independent components, in which, *ν_G,GazeG_, ν_G,GazeL_, ν_L,GazeG_* and *ν_L,GazeL_* were all set as free parameters. For all of four models, we set response bias to be unconstrained, i.e., z was a free parameter. Thus, aDDM 1 and aDDM 3 were in fact identical to DDM 2 and DDM 4; in other words, aDDM 1 and aDDM 3 were not really “attentional” DDM, as the impact of attention was constrained in the two models. Here, we treated them as reduced forms of aDDM 4, and therefore we labeled them as aDDM1 and aDDM3 for ease of comparison. The four aDDMs were fit to choice and response time data in the same way as the four DDMs except that only ‘valid’ eye-tracking trials (see above) were included for aDDM analysis.

In addition, we tested two other models that also allowed valuation bias to have both gaze-dependent and -independent components but had one more constraint compared to aDDM 4: aDDM 5, a model that allowed *ν_G,GazeG_* and *ν_G,GazeL_* to be free but set *ν_L,GazeG_* = *ν_L,GazeL_*; and aDDM 6, a model allowed *ν_L,GazeG_* and *ν_L,GazeL_* to be free but set *ν_G,GazeG_ = ν_G,GazeL_*. The performance of these two models were similar to but inferior to aDDM 4. See *SI Appendix,* Table S2 for a summary of all aDDMs.

### Prospect Theory Model

Based on the established approach (3, 9), the expected utility of accepting a gamble was calculated as the sum of probability-weighted gain and loss, i.e., U(Accept) = 0.5 × Gain – 0.5 × λ × Loss, where λ indicated relative weighting of loss over gain. The expected utility to reject a gamble is 0, i.e., U(Reject)=0. The probability of accepting a gamble was estimated based on the softmax function, P(Accept)=1/(1+e^μ-(U(Accept)-U(Reject))^), where μ indicated the degree to which a decision was sensitive to expected utility. The probability of rejecting was computed as P(Reject)=1-P(Accept). Maximum likelihood estimation was then employed to identify the values of λ and μ. The estimation was implemented using a hierarchical Bayesian approach (hBayesDM package in R) (45).

### Regression Analysis and *t*Tests

For all regression analysis, regressors were rescaled between 0 and 1. For multi-level regressions, random effects of participants were included for the intercept, and all regressors. To compare the sizes of betas between each pair of regressors, linear contrasts were performed on the absolute values of betas using the linhyptest function in MATLAB. All *t*Tests were one-tailed.

## Author contributions

F.S., A.R. and M.L.P. designed research, F.S., A.R. and P.C. performed research, F.S., A.R. S.D., W.J.Z. and S.T. analyzed data, and F.S., A.R. and M.L.P. wrote the paper.

## Data Availability

Behavioral and eye-tracking data are available at https://osf.io/yz9e6/.

## Acknowledgments

We thank the Wharton Behavioral Lab and Robert Botto for help with setting up the experiment. This work was supported by a pilot grant awarded to F.S. and A.R. from the Penn ALACRITY Center (P50-MH-113840). A.R. was supported in part by the P&S fund and a Brain and Behavior Research Foundation grant 27649. M.L.P. was supported in part by NIH grants R37-MH-109728-01, R01-MH-108627-01, Simon’s Foundation grant 304935, and the Wharton Neuroscience Initiative.

## References

1. D. Kahneman, A. Tversky, Prospect theory: An analysis of decision under risk. Econometrica 47, 263–292 (1979).

2. S. M. Tom, C. R. Fox, C. Trepel, R. A. Poldrack, The neural basis of loss aversion in decision-making under risk. Science 315, 515–518 (2007).

3. P. Sokol-Hessner, et al., Thinking like a trader selectively reduces individuals’ loss aversion. Proc. Natl. Acad. Sci. 106, 5035–5040 (2009).

4. B. De Martino, C. F. Camerer, R. Adolphs, Amygdala damage eliminates monetary loss aversion. Proc. Natl. Acad. Sci. 107, 3788–3792 (2010).

5. P. Sokol-Hessner, R. B. Rutledge, The psychological and neural basis of loss aversion. Curr. Dir. Psychol. Sci. 28, 20–27 (2019).

6. S. F. Brosnan, et al., Endowment Effects in Chimpanzees. Curr. Biol. 17, 1704–1707 (2007).

7. M. K. Chen, V. Lakshminarayanan, L. R. Santos, How Basic Are Behavioral Biases? Evidence from Capuchin Monkey Trading Behavior. J. Polit. Econ. 114, 517–537 (2006).

8. M. Bhatti, H. Jang, J. D. Kralik, J. Jeong, Rats exhibit reference-dependent choice behavior. Behav. Brain Res. 267, 26–32 (2014).

9. A. Tversky, D. Kahneman, Advances in prospect theory: Cumulative representation of uncertainty. J. Risk Uncertain. 5, 297–323 (1992).

10. B. A. Tversky, P. Slovic, D. Kahneman, The Causes of Preference Reversal. Am. Econ. Rev. 80, 204–217 (1990).

11. A. Tversky, I. Simonson, Context-Dependent Preferences. Manage. Sci. 39, 1179–1189 (1993).

12. A. Tversky, S. Sattath, P. Slovic, Contingent Weighting in Judgment and Choice. Psychol. Rev. 95, 371–384 (1988).

13. P. Slovic, S. Lichtenstein, Relative importance of probabilities and payoffs in risk taking. J. Exp. Psychol. 78, 1–18 (1968).

14. I. Ritov, J. Baron, Status-quo and Omission Biases. J. Risk Uncertain. 5, 49–61 (1992).

15. W. M. Goldstein, H. J. Einhorn, Expression Theory and the Preference Reversal Phenomena. Psychol. Rev. 94, 236–254 (1987).

16. R. Dhar, I. Simonson, The Effect of Forced Choice on Choice. J. Mark. Res. 40, 146–160 (2003).

17. E. Ert, I. Erev, The rejection of attractive gambles, loss aversion, and the lemon avoidance heuristic. J. Econ. Psychol. 29, 715–723 (2008).

18. W. Samuelson, R. Zeckhauser, Status Quo Bias in Decision Making. J. Risk Uncertain. 1, 7–59 (1988).

19. M. N. Shadlen, W. T. Newsome, Motion perception: Seeing and deciding. Proc. Natl. Acad. Sci. 93, 628–633 (1996).

20. R. Ratcliff, P. L. Smith, S. D. Brown, G. McKoon, Diffusion decision model: Current issues and history. Trends Cogn. Sci. 20, 260–281 (2016).

21. R. Polanía, I. Krajbich, M. Grueschow, C. C. Ruff, Neural oscillations and synchronization differentially support evidence accumulation in perceptual and value-based decision making. Neuron 82, 709–720 (2014).

22. A. C. Huk, M. N. Shadlen, Neural activity in macaque parietal cortex reflects temporal integration of visual motion signals during perceptual decision making. J. Neurosci. 25, 10420–10436 (2005).

23. T. D. Hanks, et al., Distinct relationships of parietal and prefrontal cortices to evidence accumulation. Nature 520, 220–223 (2015).

24. W. J. Zhao, L. Walasek, & S. Bhatia, Psychological Mechanisms of Loss Aversion: A Drift-Diffusion Decomposition. PsyArXiv:10.31234/osf.io/eg8br (28 Nov 2019).

25. S. N. Clay, J. A. Clithero, A. M. Harris, C. L. Reed, Loss aversion reflects information accumulation, not bias: A drift-diffusion model study. Front. Psychol. 8, 1–12 (2017).

26. S. Shimojo, C. Simion, E. Shimojo, C. Scheier, Gaze bias both reflects and influences preference. Nat. Neurosci. 6, 1317–1322 (2003).

27. I. Krajbich, C. Armel, A. Rangel, Visual fixations and the computation and comparison of value in simple choice. Nat. Neurosci. 13, 1292–1298 (2010).

28. J. F. Cavanagh, T. V. Wiecki, A. Kochar, M. J. Frank, Eye tracking and pupillometry are indicators of dissociable latent decision processes. J. Exp. Psychol. Gen. 143, 1476–1488 (2014).

29. I. Krajbich, D. Lu, C. Camerer, A. Rangel, The attentional drift-diffusion model extends to simple purchasing decisions. Front. Psychol. 3 (2012).

30. S. Joshi, Y. Li, R. M. Kalwani, J. I. Gold, Relationships between pupil diameter and neuronal activity in the locus coeruleus, colliculi, and cingulate cortex. Neuron 89, 221–234 (2016).

31. S. Mathôt, Pupillometry: Psychology, Physiology, and Function. J. Cogn. 1, 1–23 (2018).

32. M. A. Just, P. A. Carpenter, A. Miyake, Neuroindices of cognitive workload: Neuroimaging, pupillometric and event-related potential studies of brain work. Theor. Issues Ergon. Sci. 4, 56–88 (2003).

33. J. W. de Gee, T. Knapen, T. H. Donner, Decision-related pupil dilation reflects upcoming choice and individual bias. Proc. Natl. Acad. Sci. 111, E618–E625 (2014).

34. E. Ert, I. Erev, On the descriptive value of loss aversion in decisions under risk: Six clarifications. Judgm. Decis. Mak. 8, 214–235 (2013).

35. E. Yechiam, Acceptable losses: the debatable origins of loss aversion. Psychol. Res. 83, 1327–1339 (2019).

36. T. Pachur, M. Schulte-mecklenbeck, R. O. Murphy, R. Hertwig, Prospect theory reflects selective allocation of attention. J. Exp. Psychol. Gen. 147, 147–169 (2018).

37. B. Hoeks, W. J. M. Levelt, Pupillary dilation as a measure of attention: a quantitative system analysis. Behav. Res. Methods, Instruments, Comput. 25, 16–26 (1993).

38. N. Canessa, et al., The functional and structural neural basis of individual differences in loss aversion. J. Neurosci. 33, 14307–14317 (2013).

39. P. Sokol-Hessner, et al., Determinants of propranolol’s selective effect on loss aversion. Psychol. Sci. 26, 1123–1130 (2015).

40. H. Takahashi, et al., Norepinephrine in the brain is associated with aversion to financial loss. Mol. Psychiatry 18, 3–4 (2013).

41. M. E. Kret, E. E. Sjak-Shie, Preprocessing pupil size data: Guidelines and code. Behav. Res. Methods 51, 1–7 (2018).

42. T. V. Wiecki, I. Sofer, M. J. Frank, HDDM: Hierarchical Bayesian estimation of the Drift-Diffusion Model in Python. Front. Neuroinform. 7, 1–10 (2013).

43. A. Gelman, D. B. Rubin, Inference from Iterative Simulation Using Multiple Sequences. Stat. Sci. 7, 457–472 (1992).

44. T. Ando, Bayesian predictive information criterion for the evaluation of hierarchical Bayesian and empirical Bayes models. Biometrika 94, 443–458 (2007).

45. W.-Y. Ahn, N. Haines, L. Zhang, Revealing neurocomputational mechanisms of reinforcement learning and decision-making with the hBayesDM package. Comput. Psychiatry 1, 24–57 (2017).

