## Supplementary material for "Decomposing Loss Aversion from Gaze Allocation and Pupil Dilation": SI Appendix

**This PDF file includes:**

Supplementary text

Figures S1 to S8

Tables S1 to S2

**Supplementary text**

According to the attentional drift diffusion model (aDDM), gaze on one option or feature will discount the weight of the nonfixated option or feature in drift rate. If this is the case, valuation bias, or the relative weight of loss vs. gain in drift rate, $v_L/v_G$, should be larger when gaze is on loss than when gaze is on gain. To directly test this idea, we developed another DDM that incorporated gaze-loss ratio and allowed the coefficients for gain and loss in the drift rate to vary as a function of gaze, i.e., $v_{G,GazeG}$ and $v_{L,GazeG}$ for gaze on gain and $v_{G,GazeL}$ and $v_{L,GazeL}$ for gaze on loss (*Materials and Methods*). We found this model outperformed other models that had constraints on $v_{G,GazeG}$, $v_{L,GazeG}$, $v_{G,GazeL}$ and $v_{L,GazeL}$ (Fig. S8*A* and Table S2). Importantly, based on this model, we found valuation bias was larger when gaze was on loss (Fig. S8*B*, dark brown bar, $v_{L,GazeL}/v_{G,GazeL}$=2.022, 95% credible interval: 1.682 to 2.486) than when gaze was on gain (light brown bar, $v_{L,GazeG}/v_{G,GazeG}$=1.241, 95% credible interval: 1.096 to 1.405, greater than 99.9% likelihood $v_{L,GazeL}/v_{G,GazeL} > v_{L,GazeG}/v_{G,GazeG}$). This effect was mainly driven by the discounted coefficient of gain when gaze was on loss relative to when gaze was on gain ($v_{G,GazeL}/v_{G,GazeG}$=0.647, 95% credible interval: 0.511 to 0.800, greater than 99.9% likelihood < 1), while the coefficient of loss was stable regardless of whether gaze was on gain or loss ($v_{L,GazeG}/v_{L,GazeL}$=0.949, 95% credible interval: 0.819 to 1.094, 75.8% likelihood < 1). See Table S2 for details.

These findings unveil the dynamic relationship between gaze allocation and valuation bias in the evidence accumulation process. However, this does not mean that valuation bias is driven solely by gaze bias. It is notable that even when gaze was on gain, valuation bias of weighting loss over gain was significant (Fig. S8*B*, light brown bar, $v_{L,GazeG}/v_{G,GazeG}$, greater than

99.9% likelihood > 1). In fact, when we held this aDDM model as unconstrained and compared it with two constrained models, the one that assumed valuation bias completely depended on gaze (i.e., $v_{G,GazeG}=v_{L,GazeL}$ and $v_{G,GazeL}=v_{L,GazeG}$, Fig. S8*A*, orange bar) explained less variance than the model that assumed valuation bias was completely independent of gaze (i.e., $v_{G,GazeG}=v_{G,GazeL}$ and $v_{L,GazeG}=v_{L,GazeL}$, Fig. S8*A*, gray bar), though both of the two constrained models underperformed the unconstrained model, which allowed valuation bias to have both gaze-dependent and gaze-independent components (Fig. S8*A*, brown bar; see Table S2 for details). Based on these findings, a plausible mechanism is that loss-averse decision makers have an initial *a priori* inclination to weigh loss over gain as evidence supporting a decision, which drives them to preferentially inspect information about loss relative to gain with gaze, and this gaze bias, in turn, further enhances the weight of loss in the evidence accumulation process.

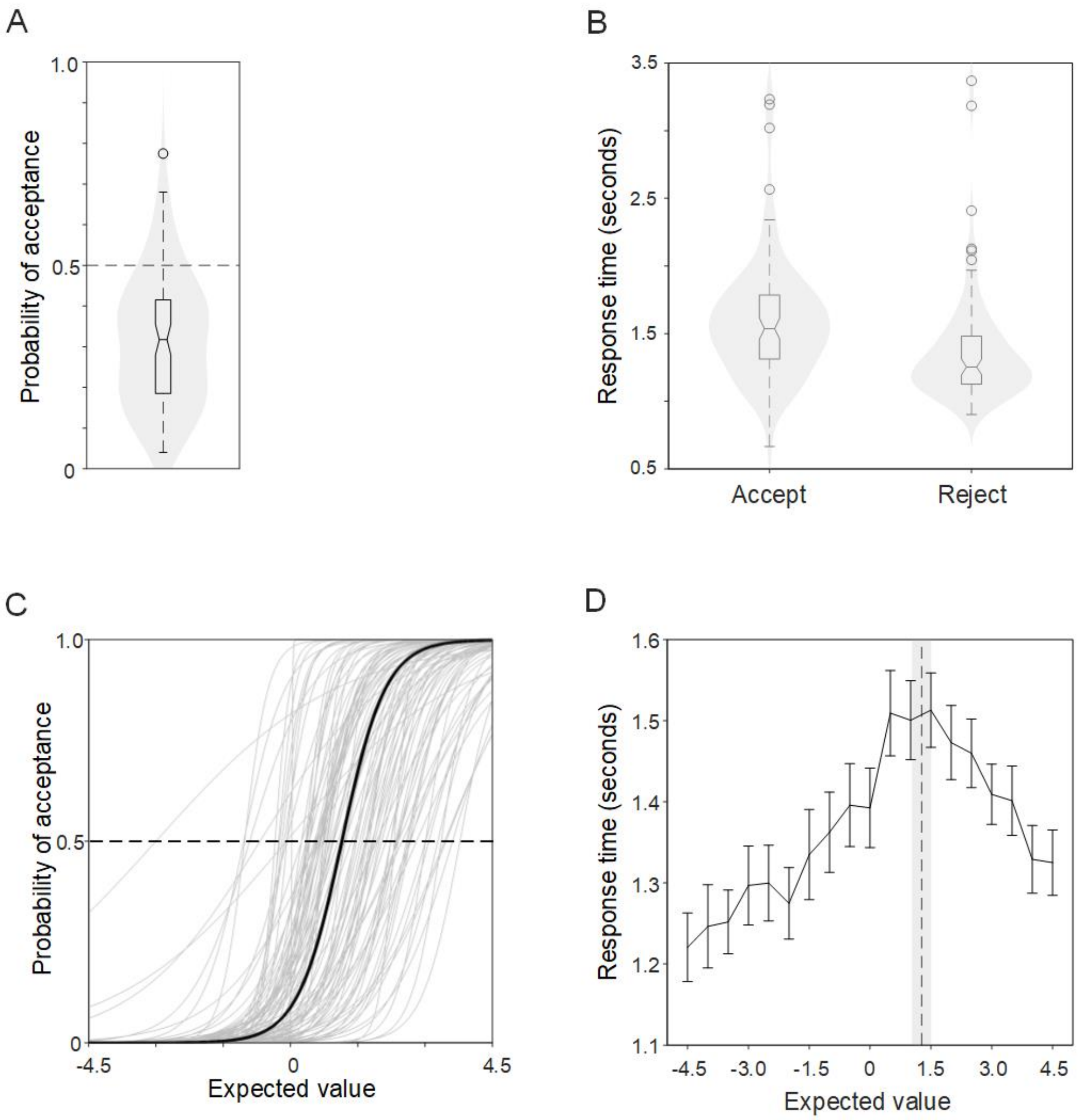


**Fig. S1.** Choices and response times. (*A*) Violin plot of probability of gamble acceptance across participants. (*B*) Violin plot of response times for accepting and rejecting gambles across participants. (*C*) Probability of gamble acceptance conditional on expected values (grey curves and black curves were fit with logistic regressions across individuals and the group, respectively). (*D*) Response times conditional on expected values. Error bars indicate SEs. The dashed black line indicates the mean expected value that evoked longest response times of the group, and the shaded area indicates standard error.

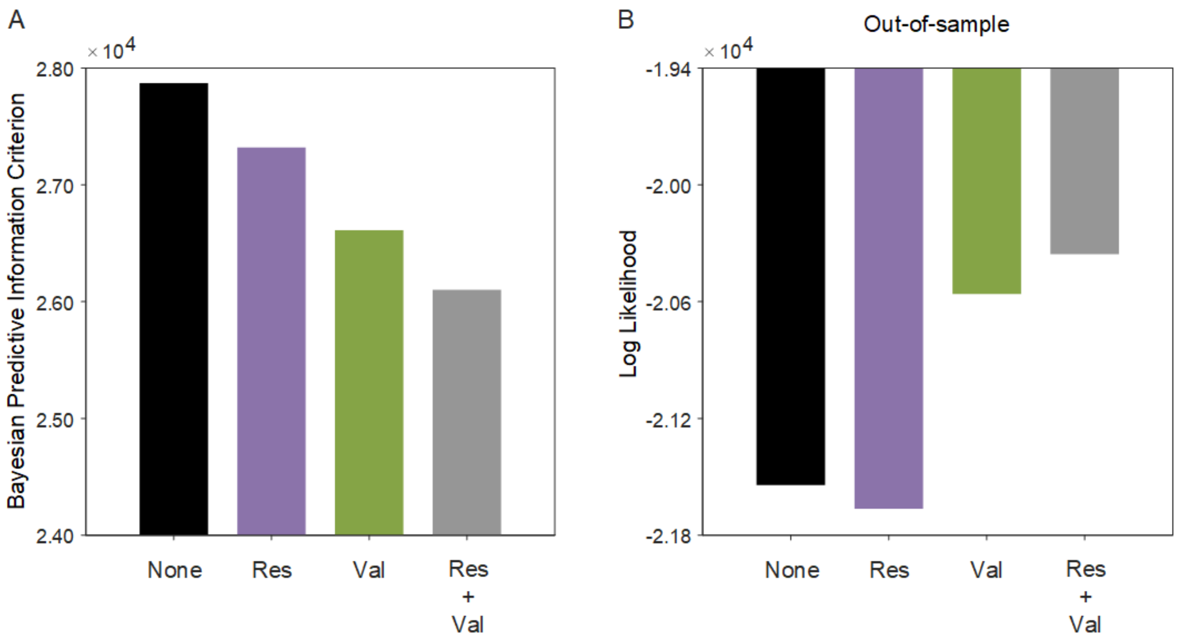


**Fig. S2.** Model fit. (*A*) Bayesian Predictive Information Criterion (BPIC) for the four models. A model with a smaller BPIC score explained more variance in choice probabilities and response times. (*B*) Log likelihood based on out-of-sample prediction that used posterior means of parameters estimated with choice and response time data of gain-left (gain-right) trials to predict the joint probability distribution of choice and response time of gain-right (gain-left) trials. Res: Response bias. Val: Valuation bias.

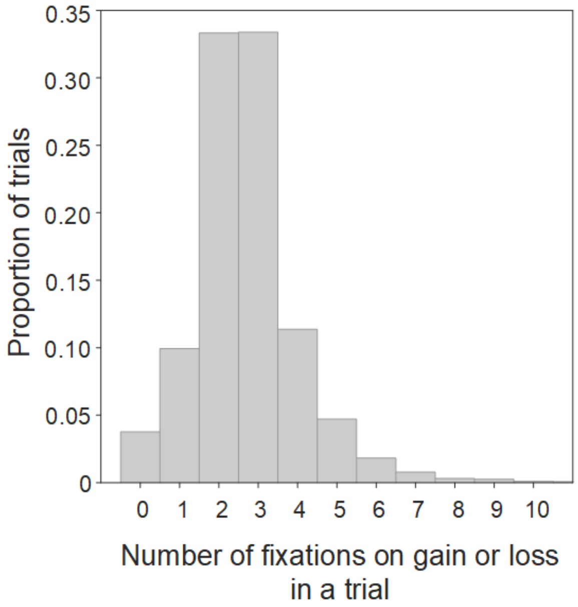


**Fig. S3.** Gaze fixations. Histogram of the number of gaze fixations on gain and loss amounts across trials. Trials with more than 10 fixations on gain or loss were rare (0.002) and not illustrated in the figure.

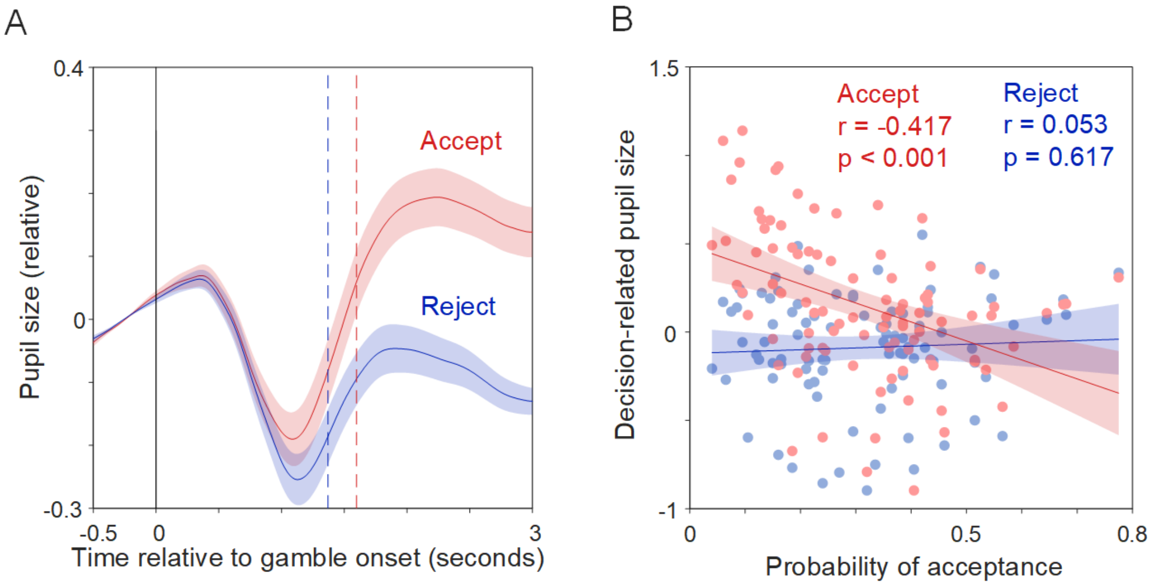


**Fig. S4.** Pupil dilation conditional on decisions. (*A*) The time course of pupil size aligned to gamble onset. The dashed vertical lines indicated mean response times for gamble acceptance (red) and rejection (blue). (*B*) Correlation between probability of gamble acceptance and pupil size separated by accept decisions (red) and reject decisions (blue). Each participant is represented by a red dot and a blue dot.

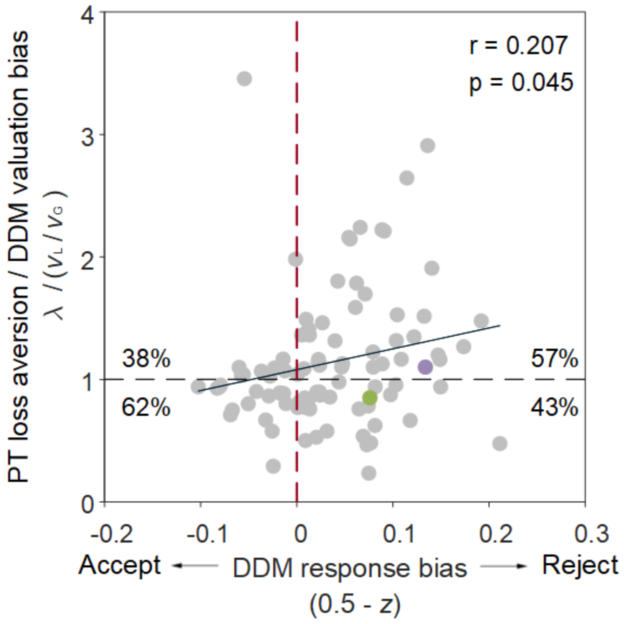


**Fig. S5.** The ratio of loss aversion estimated by prospect theory (PT) to valuation bias estimated by DDM with respect to response bias. The green dot (P69, quadrant 4) and the purple dot (P46, quadrant 1) indicate the two example participants illustrated in **Fig. S6** and **Fig. S7**.

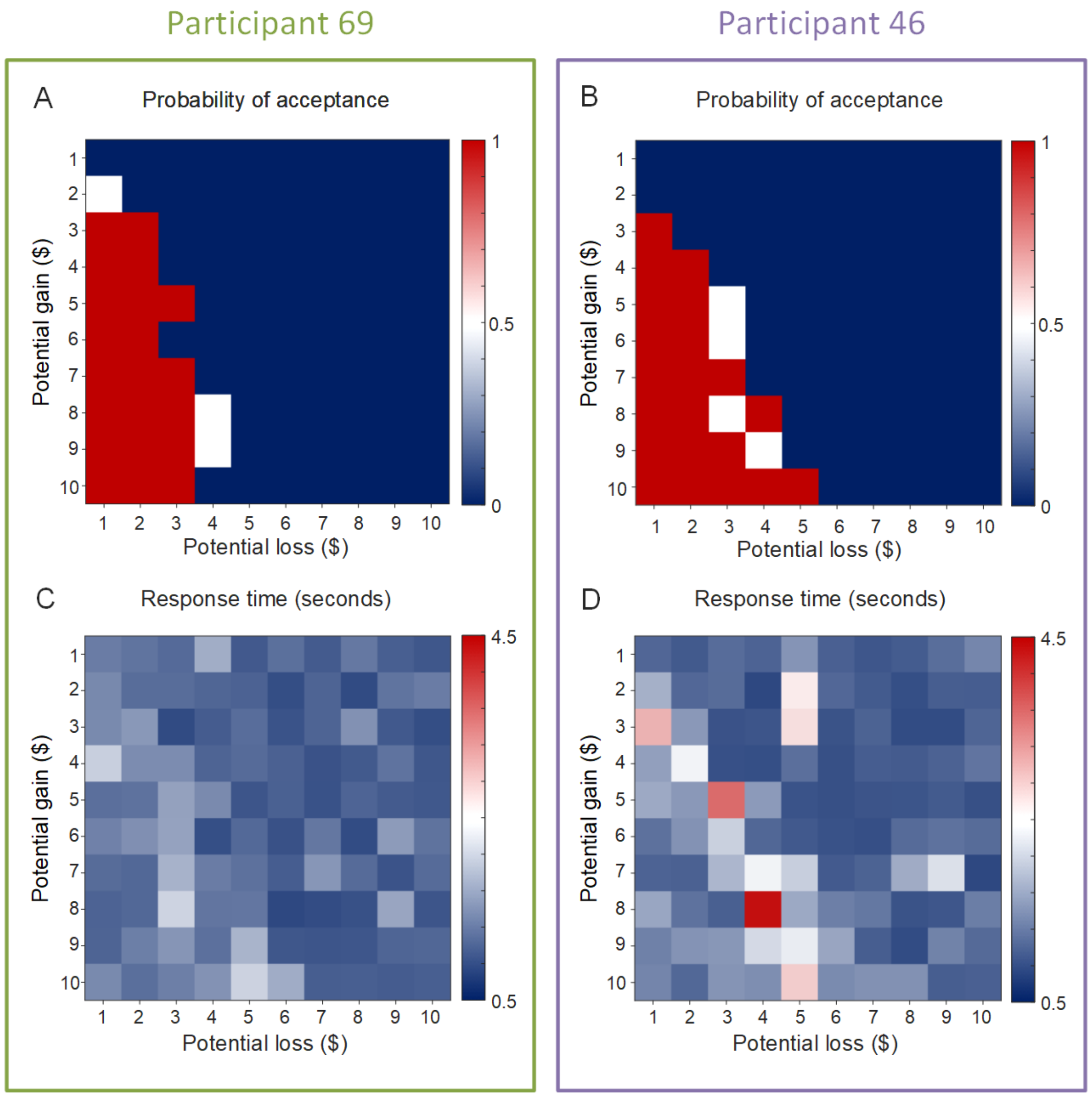


**Fig. S6.** Example participants' choices and response times. Probability of acceptance (*A*, *B*) and response times (*C*, *D*) of two example participants who are denoted by the colored dots (green and purple) in **Fig. 1E** and **Fig. S5.**

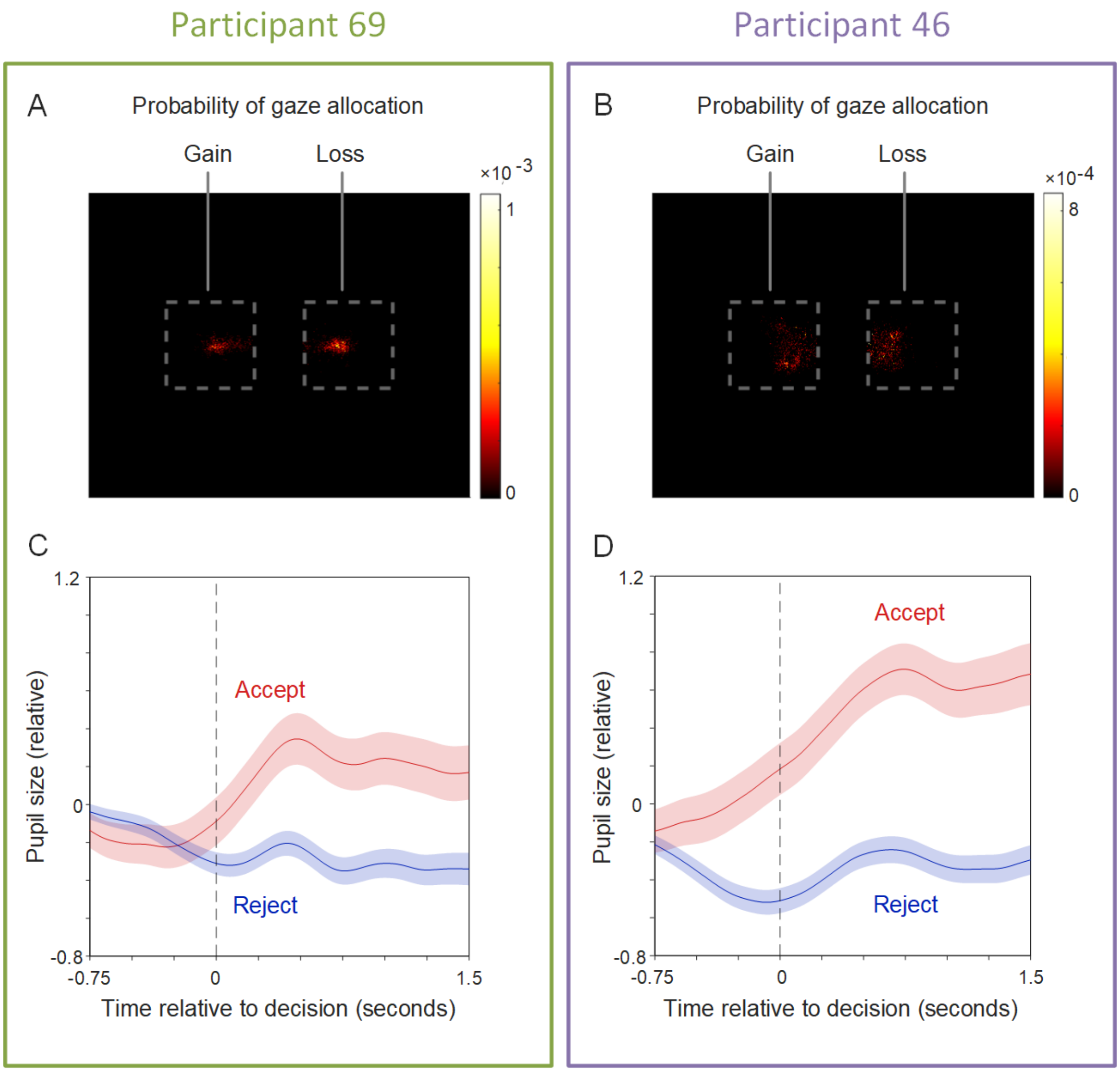


**Fig. S7.** Example participants' gaze allocation and pupil dilation. Gaze allocation (*A*, *B*) and pupil dilation (*C*, *D*) of two example participants who are denoted by the colored dots (green and purple) in **Fig. 1E** and **Fig. S5.**

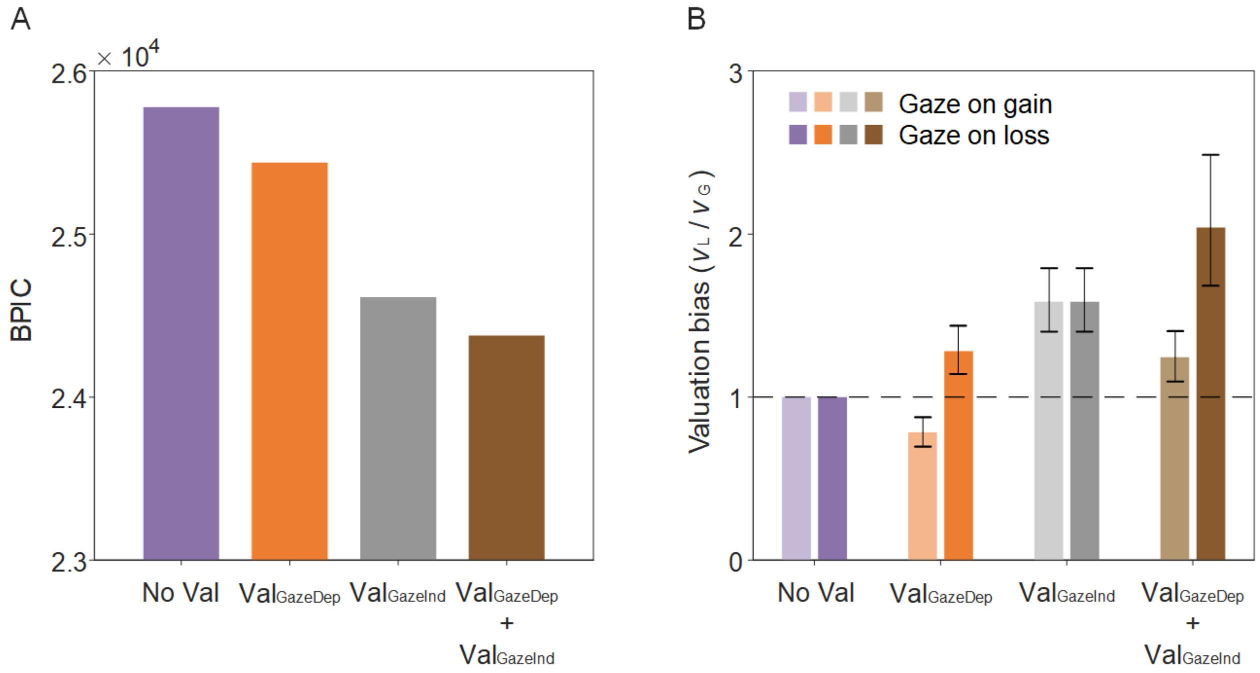


**Fig. S8.** aDDM. (*A*) Bayesian Predictive Information Criterion (BPIC) and (*B*) valuation bias estimated from different aDDMs. Error bars indicate 95% credible intervals. Val: Valuation bias. $Val_{GazeDep}$: Gaze-dependent valuation bias. $Val_{GazeInd}$: Gaze-independent valuation bias. See Table S2 for details.

**Table S1.** Summary of DDM estimates (mean and 95% credible interval). Res: Response bias; Val: Valuation bias.

| Index | Color | Allowed bias(es) | Parameters | BPIC | Bayesian estimates of the group | | | Valuation bias | Response bias |
|---|---|---|---|---|---|---|---|---|---|
| | | | | | $v_G$ | $v_L$ | $z$ | $v_L / v_G$ | $0.5 - z$ |
| DDM 1 | | - | $z = 0.5$; $v_G = v_L$ | 27871 | 0.271 (0.251, 0.290) | 0.271 (0.251, 0.290) | 0.5 | 1 | 0 |
| DDM 2 | | Res | $z$; $v_G = v_L$ | 27321 | 0.272 (0.253, 0.292) | 0.272 (0.253, 0.292) | 0.463 (0.448, 0.477) | 1 | 0.037 (0.023, 0.052) |
| DDM 3 | | Val | $z = 0.5$; $v_G$; $v_L$ | 26612 | 0.216 (0.194, 0.238) | 0.339 (0.315, 0.364) | 0.5 | 1.569 (1.391, 1.779) | 0 |
| DDM 4 | | Res + Val | $z$; $v_G$; $v_L$ | 26100 | 0.218 (0.196, 0.239) | 0.342 (0.318, 0.366) | 0.463 (0.448, 0.477) | 1.571 (1.398, 1.780) | 0.037 (0.023, 0.052) |

**Table S2.** Summary of aDDM estimates (mean and 95% credible interval). Res: Response bias; $\text{Val}_{\text{GazeDep}}$: Gaze-dependent valuation bias; $\text{Val}_{\text{GazeInd}}$: Gaze-independent valuation bias.

| Index | Color | Allowed bias(es) | Parameters | BPIC | Bayesian estimates of the group | | | | Valuation bias | |
|---|---|---|---|---|---|---|---|---|---|---|
| | | | | | $v_{\text{G, GazeG}}$ | $v_{\text{G, GazeL}}$ | $v_{\text{L, GazeG}}$ | $v_{\text{L, GazeL}}$ | $v_{\text{L, GazeG}}$ / $v_{\text{G, GazeG}}$ | $v_{\text{L, GazeL}}$ / $v_{\text{G, GazeL}}$ |
| aDDM 1 (DDM 2) | | Res | $v_{\text{G, GazeG}} = v_{\text{L, GazeG}} = v_{\text{L, GazeG}} = v_{\text{L, GazeL}}$ | 25778 | 0.272 (0.252, 0.291) | 0.272 (0.252, 0.291) | 0.272 (0.252, 0.291) | 0.272 (0.252, 0.291) | 1 | 1 |
| aDDM 2 | | Res $\text{Val}_{\text{GazeDep}}$ | $v_{\text{G, GazeG}} = v_{\text{L, GazeL}}$ ; $v_{\text{G, GazeL}} = v_{\text{L, GazeG}}$ | 25438 | 0.300 (0.282, 0.319) | 0.235 (0.212, 0.258) | 0.235 (0.212, 0.258) | 0.300 (0.282, 0.319) | 0.782 (0.696, 0.876) | 1.278 (1.141, 1.437) |
| aDDM 3 (DDM 4) | | Res $\text{Val}_{\text{GazeInd}}$ | $v_{\text{G, GazeG}} = v_{\text{G, GazeL}}$ ; $v_{\text{L, GazeG}} = v_{\text{L, GazeL}}$ | 24613 | 0.216 (0.194, 0.238) | 0.216 (0.194, 0.238) | 0.342 (0.318, 0.366) | 0.342 (0.318, 0.366) | 1.581 (1.402, 1.791) | 1.581 (1.402, 1.791) |
| aDDM 4 | | Res $\text{Val}_{\text{GazeDep}}$ $\text{Val}_{\text{GazeInd}}$ | $v_{\text{G, GazeG}}$ ; $v_{\text{L, GazeG}}$ ; $v_{\text{L, GazeG}}$ ; $v_{\text{L, GazeL}}$ | 24377 | 0.265 (0.240, 0.292) | 0.172 (0.140, 0.204) | 0.329 (0.298, 0.362) | 0.347 (0.315, 0.380) | 1.241 (1.096, 1.405) | 2.022 (1.682, 2.486) |
| aDDM 5 | - | Res $\text{Val}_{\text{GazeDep}}$ $\text{Val}_{\text{GazeInd}}$ | $v_{\text{G, GazeG}}$ ; $v_{\text{L, GazeG}}$ ; $v_{\text{L, GazeG}} = v_{\text{L, GazeL}}$ | 24419 | 0.270 (0.249, 0.291) | 0.165 (0.137, 0.193) | 0.337 (0.314, 0.362) | 0.337 (0.314, 0.362) | 1.249 (1.126, 1.388) | 2.046 (1.716, 2.497) |
| aDDM 6 | - | Res $\text{Val}_{\text{GazeDep}}$ $\text{Val}_{\text{GazeInd}}$ | $v_{\text{G, GazeG}} = v_{\text{L, GazeG}}$ ; $v_{\text{L, GazeG}}$ ; $v_{\text{L, GazeL}}$ | 24450 | 0.216 (0.194, 0.237) | 0.216 (0.194, 0.237) | 0.291 (0.260, 0.321) | 0.384 (0.356, 0.412) | 1.349 (1.173, 1.557) | 1.780 (1.579, 2.019) |
